# The microbial composition and functional roles of different kombucha products in Singapore

**DOI:** 10.1101/2022.11.14.516366

**Authors:** Suet Li Hooi, Jacky Dwiyanto, Kai Yee Toh, Gwendoline Tan, Chun Wie Chong, Jonathan Wei Jie Lee, Jeremy Lim

## Abstract

Kombucha is a fermented tea traditionally known for its health-enhancing properties owing to the bioactive compounds generated by acetic acid bacteria (AAB) and lactic acid bacteria (LAB). We compared the distribution of AAB and LAB and their functional pathways across nine commercial kombucha products in Singapore using shotgun metagenomic sequencing. A high prevalence of *Komagataeibacter* species including *Komagataeibacter saccharivorans* (82.93% in B), *Komagataeibacter xylinus* (93.38% in D) and *Komagataeibacter rhaeticus* (92.20% and 30.62% in G and I) was detected in AAB-dominant kombucha. LAB-dominant kombucha was largely represented by *Bacillus coagulans* (~99% in E and F) and *Lactobacillus nagelii* (~60% in H). Despite differences in bacterial composition, both LAB-and AAB-dominant kombucha harbour pathways involved in the biosynthesis of short-chain fatty acids (SCFAs), amino and organic acids and vitamin B12. “Fatty acid and beta-oxidation II (peroxisome)” and “fatty acid and beta-oxidation I” were detected in the LAB but not the AAB-dominant kombucha.

## Introduction

Kombucha is a traditionally fermented drink produced from the fermentation of sweetened green, black or oolong tea by the symbiotic culture of bacteria and yeast (SCOBY) [1,2]. This traditional beverage has been widely consumed worldwide over the last decade due to its purported health benefits. For instance, regular consumption of kombucha is believed to reduce blood pressure, aid in weight loss, improve immunity, facilitate digestion, prevent the development of metabolic and gastrointestinal diseases as well as cancer [1,3]. These positive health effects of kombucha are mainly attributed to the unique microbial consortium and the bioactive compounds these microbes produce during the fermentation process [4].

The SCOBY in kombucha is primarily made up of yeast and a large proportion of bacteria called acetic acid bacteria (AAB). AAB are Gram-negative obligate aerobes (which strictly require oxygen to survive) belonging to the *Acetobacteraceae* family. They can grow optimally at temperatures of approximately 25 to 30 °C and a pH of 5.0 to 6.5 [5,6]. The most prevalent AAB in kombucha are the species of the genera *Komagataeibacter*, *Gluconobacter, Acetobacter* and *Gluconacetobacter* [6]. Common AAB species found in commercial kombucha are *Komagataeibacter xylinus*, *Komagataeibacter saccharivorans*, *Acetobacter xylinum* and *Gluconobacter oxydans* [3,7,8]. These bacteria produce acetic acid and cellulose through the fermentation process. The metabolic products, in-turn confer various health-promoting properties such as antimicrobial activities and the prevention of metabolic diseases like diabetes and obesity by decreasing intestinal glucose absorption and converting glucose into cellulose [2,9].

Lactic acid bacteria (LAB) are less frequently detected in kombucha. When present, LAB are usually found in relatively low abundances [6,7,8]. However, LAB such as *Bacillus coagulans* and *Lactobacillus* species are often added in commercial kombucha fermentation to enhance the probiotic potential of the beverage [8,10]. LAB are Gram-positive, facultative anaerobic bacteria (can survive in the presence or absence of oxygen) belonging to the *Firmicutes* phylum, which grows optimally at temperatures between 25 to 40°C and a pH of 4.0 to 6.0. LAB can be categorised into either homofermenters or heterofermenters [11,12]. *Lactobacillus, Lactococcus* and *Streptococcus* species are among the most common homofermenters which utilise glucose to produce lactic acid as the primary end-product of glycolysis. Meanwhile, heterofermenters such as *Leuconostoc* and *Weisella* spp. generate lactic acid, carbon dioxide and ethanol via the pentose phosphate pathway [6,11,12]. Similar to AAB, the metabolites produced by LAB facilitate digestion, prevent infections and exert immunomodulation properties [11,12].

The different preparation and brewing processes have resulted in variable microbial composition across kombucha. In this study, we profiled the microbial composition and functional pathways of nine commercial kombucha products in Singapore using shotgun metagenomics sequencing. We further compared the microbial and functional differences between the AAB-and LAB-dominant.

## Methods

### Sample procurement and processing

Nine kombucha products were obtained from commercial supermarkets in Singapore. Each product was brewed according to the manufacturer’s instruction. The products were stored at 4°C, with a shelf-life of up to 12 months. Unpasteurised kombucha products were procured and sent to the laboratory for subsequent steps on the same day.

### DNA extraction and shotgun metagenomic sequencing

A total of 300 to 350mL of each product was centrifuged at 6000×g. The resulting supernatant was removed, and the pellet was stored at −80°C. The DNA from the frozen pellet was extracted using QIAamp PowerFecal Pro DNA kit according to the manufacturer’s protocol and quantified using Qubit 4 fluorometer. DNA sequencing was performed using Illumina NovaSeq with 2×150bp paired-end configuration and a read depth of 20 million paired-end reads. The raw sequence reads were processed using BioBakery3 workflows [13]. Read-level quality control was conducted using KneadData v0.11.0 with default settings [13]. Taxonomy annotations were performed using MetaPhlan version 3.0 which uses clade-specific markers to identify microbial species and their relative abundances. Meanwhile, functional pathways annotations were conducted using HUMAnN version 3.0 based on the MetaCyc pathway database [13].

## Data analysis

Species and pathway abundances were analysed using phyloseq R package version 1.36.0 [14]. Species and functional pathways with <10% and <0.01% relative abundances respectively were discarded using genefilter_sample and prune_taxa functions from the phyloseq R package version 1.36.0 [14]. The bar plots were generated using ggplot2 R package version 3.3.5 whereas the heatmap was generated using ComplexHeatmap R package version 2.13.1 [15,16]. The products were categorised as AAB- or LAB-dominant if the relative abundances of either AAB or LAB species reached more than 30%. Non-AAB- and LAB-dominant products (relative abundances of AAB, LAB or other species (<30%) were categorised as “others”.

## Results

All nine products were unpasteurised, and most (*n* = 5) used either SCOBY or acidifier (*n* = 2) as the starter culture. *Bacillus coagulans* which was added in E and F and was detected in our analysis with relative abundances of almost 100% (99.95% and 99.98% respectively). However, the species of *Lactobacillus*, *Acetobacter*, *Gluconobacter* and *Saccharomyces,* which claimed to be added in products D and G by the respective manufacturers were either below detection limit or present in low abundance (2.06% of *Saccharomyces cerevisiae* in G) **(Table 1** and **Fig 1a).**

**Table 1.** Commercial claims and microbiome profiles of the kombucha products (*n* = 9) included in this study.

| **Starter culture** | **Product** | **Type of dominant bacteria present** | **Probiotics added\*** | **Number of microbial species** | **Number of functional pathways** | **Average pathways per species** |
| --- | --- | --- | --- | --- | --- | --- |
| Acidifier | A | Others | None | 27 | 314 | 11.63 |
| Acidifier | B | AAB | None | 4 | 130 | 32.50 |
| SCOBY | C | Others | None | 96 | 390 | 4.06 |
| SCOBY | D | AAB | Yeast and *Lactobacillus* species | 4 | 148 | 37 |
| Unknown | E | LAB | *Bacillus coagulans* | 3 | 117 | 58.5 |
| Unknown | F | LAB | *Bacillus coagulans* | 2 | 137 | 45.67 |
| SCOBY | G | AAB | Yeast of the *Saccharomyces* genus and bacteria from *Acetobacter* and *Gluconobacter* species | 3 | 97 | 32.33 |
| SCOBY | H | LAB | None | 7 | 192 | 27.43 |
| SCOBY | I | AAB | None | 6 | 142 | 23.67 |

\* As claimed by the kombucha manufacturer

**Fig 1a.**
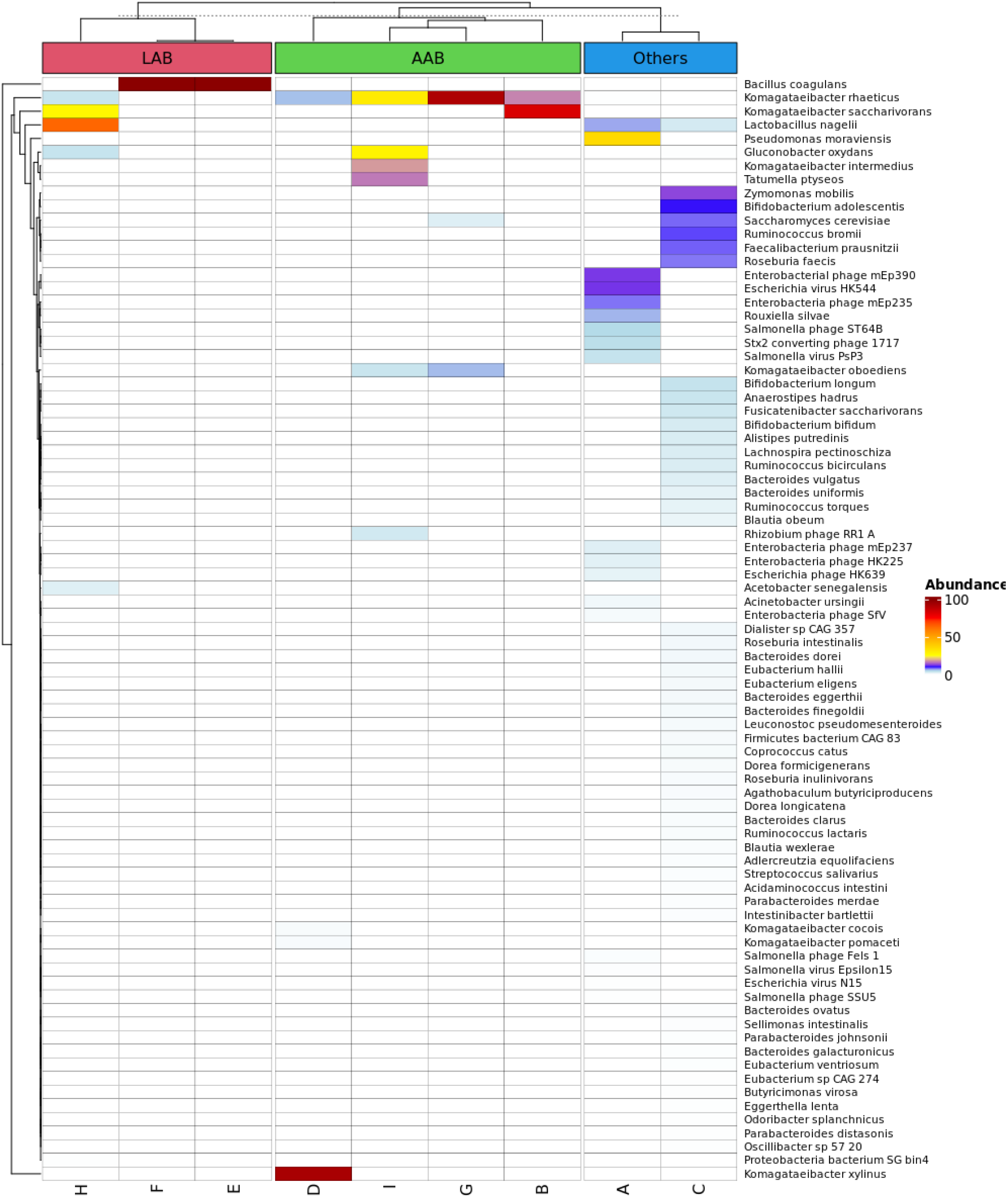
Heatmap of microbial species. The heatmap of 81 microbial species detected across the nine kombucha products after filtering for species with >10% relative abundance.

Overall, 141 microbial species were detected across all nine products. Eighty-one species remained after filtering for low species abundance (<10%) **(Fig 1a)**. Product C had the highest species diversity, followed by product A and the remaining products contained less than ten species **(Table 1)**.

The classification of species from AAB and LAB was based on the definition set by previous literature [5,11]. Four out of the nine products were predominated by AAB, notably the species from the *Komagataeibacter* genus (relative abundance > 30%). These four products, B, D, G and I were predominated by *Komagataeibacter saccharivorans* (B), *Komagataeibacter xylinus* (D) and *Komagataeibacter rhaeticus* (G and I) with relative abundances of 82.93%, 93.38%, 92.20% and 30.62%, respectively. LAB-dominant products were E, F and H, with relative abundances of *Bacillus coagulans* in E and F at 99.95% and 99.98%, respectively and *Lactobacillus nagelli* in H at 64.12% **(Fig 1a)**.

After removing functional pathways with <0.01% relative abundance, 59 out of 433 pathways remained. The top five most abundant functional pathways were the pentose phosphate pathway (non-oxidative branch), aerobic respiration I (cytochrome c), aerobic respiration II (cytochrome c) (yeast), guanosine deoxyribonucleotides de novo biosynthesis II and adenosine deoxyribonucleotides de novo biosynthesis II **(Fig 2a)**.

**Fig 2a.**
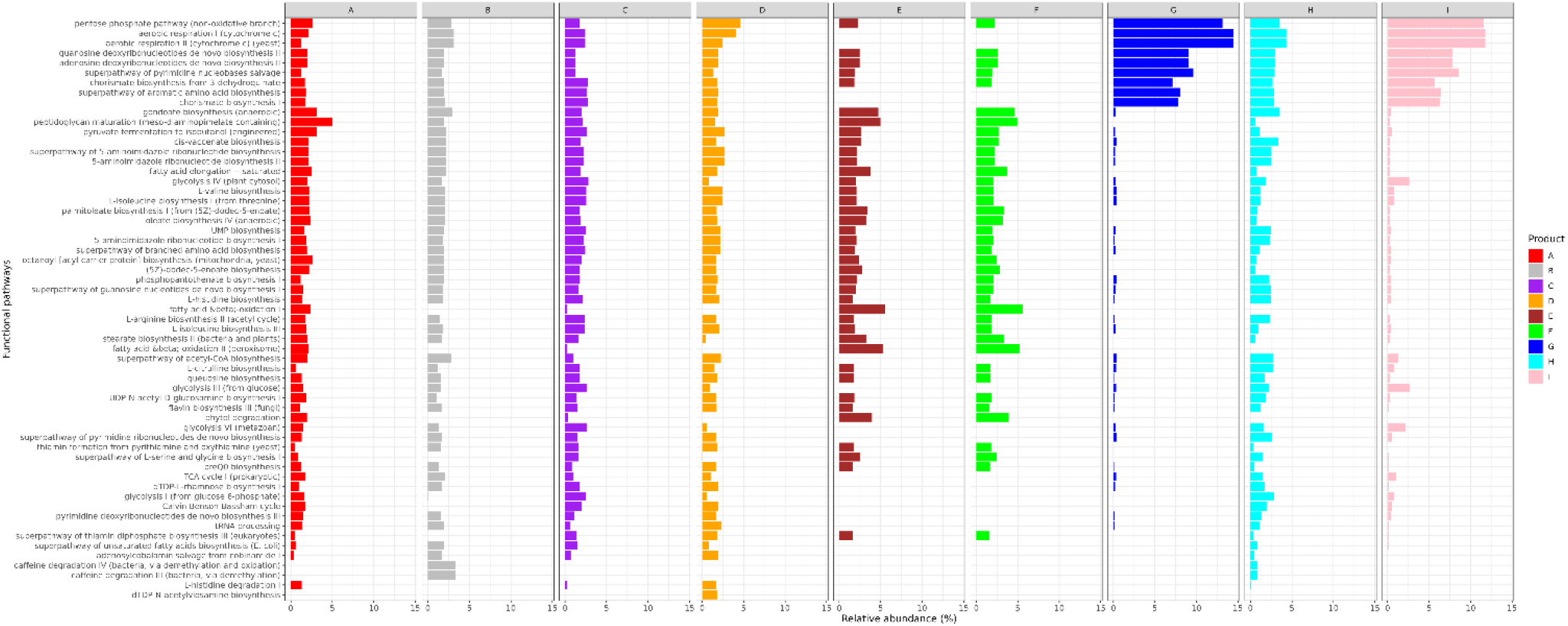
Functional pathways. The top 59 functional pathways after removing those with less than 0.01% relative abundance (the relative abundances were normalised to 100% in this figure) across all the nine kombucha products.

Of the 59 pathways, we further selected 20 that are involved in the production of beneficial metabolites. AAB was found to harbour 15 out of the 20 selected pathways (75%), while 11 pathways (55%) were detected in LAB. Notably, 2 out of 20 functional pathways (10%) were exclusively contributed by LAB and “others”. These pathways were “fatty acid and beta-oxidation II (peroxisome)” and “fatty acid and beta-oxidation I”. Meanwhile, 6 out of 20 (30%) were exclusively contributed by AAB and “others” **(Fig 2b)**.

**Fig 2b.**
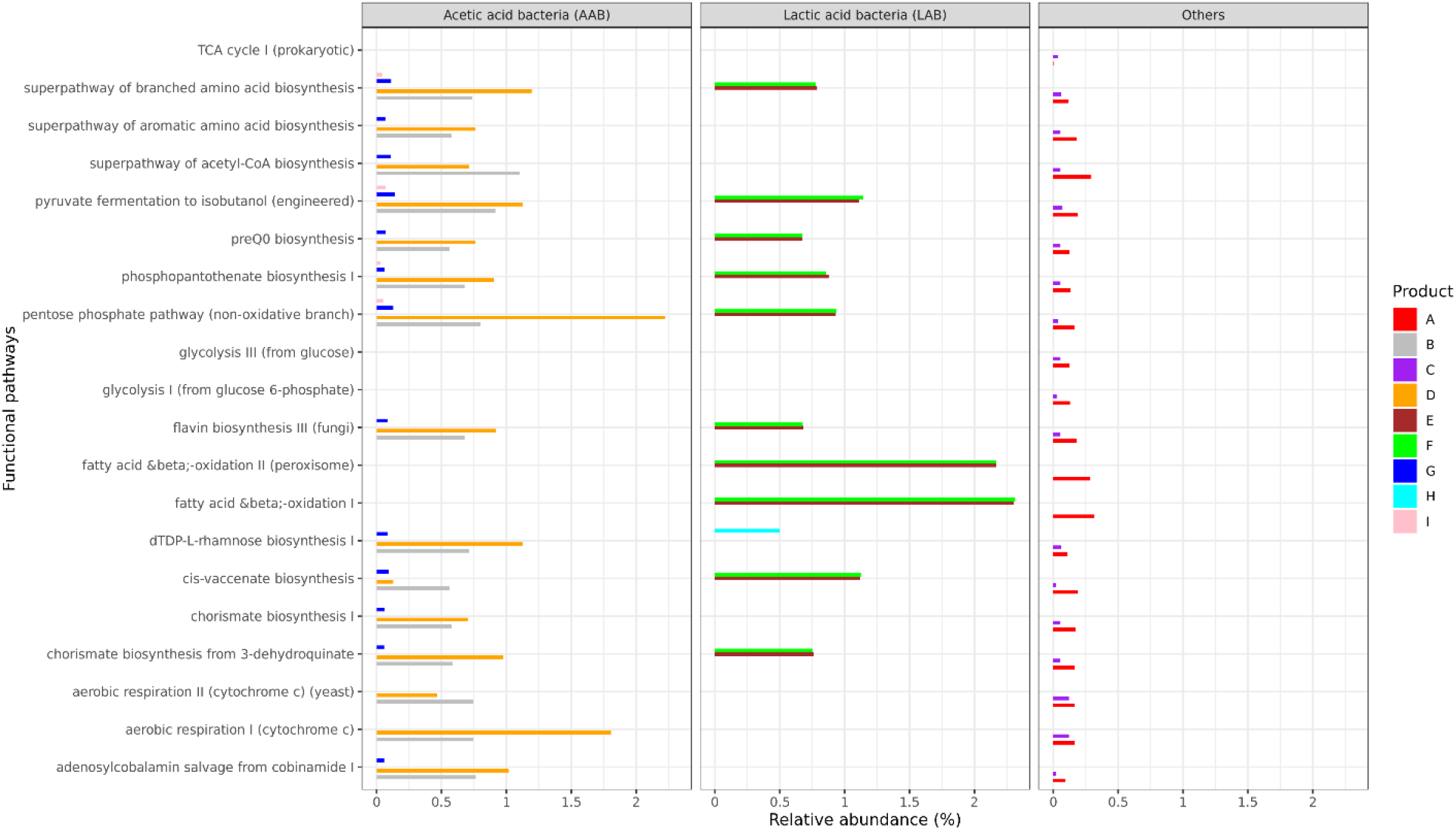
Species contribution. The relative abundances of the top 20 functional pathways contributed by three categories of microbes; AAB, LAB and others. Unclassified species contributing to the pathways across the nine products were excluded.

## Discussion

The growing popularity of kombucha in the last decade has motivated many beverage companies to introduce various new fermentation strategies, starter cultures and additional flavours, which results in unique microbial and metabolite profiles among different kombucha products. Notably, the bacterial composition in kombucha is comprised mainly of AAB and LAB. These bacteria play an essential role in producing bioactive compounds, thereby contributing to the pro-health effects of kombucha [3,6]. In this study, we analysed nine commercial kombucha products in Singapore and compared the composition of AAB and LAB and their functional potentials. Shotgun metagenomics sequencing revealed that the studied kombucha can be largely classified into three categories, namely the AAB- dominant (B, D, G and I), LAB-dominant (E, F and H) and others (A and C).

Consistent with other parallel kombucha studies, *Komagateibacter saccharivorans*, *Komagataeibacter xylinus* and *Komagataeibacter rhaeticus* are the most abundant and prevalent *Komagataeibacter* species found in the AAB-dominant products [8,17,18]. Interestingly, the high prevalence of *Komagataeibacter* spp. was found across geographically distant locations (e.g. Australia, China, Turkey and USA), suggesting a degree of standardisation in the kombucha preparation [7–8,17–19]. The species of *Komagataeibacter* are reported to exhibit efficient glucose-metabolising activities (e.g. high glucose conversion rate to cellulose) which have a putative role in preventing the onset of metabolic diseases such as type-2 diabetes and obesity [9,18].

Our study found the three LAB-dominant products (E, F and H) were predominated by *Bacillus coagulans* (~99% relative abundances in E and F) and *Lactobacillus nagelii* (64.12% relative abundance in H). Our finding was consistent with the previous studies conducted in America which also detected the predominance of these LAB species in their tested kombucha samples [7,8]. *B. coagulans* are often added to food products not only due to their numerous probiotic characteristics but also their high tolerance to extreme environments such as high temperature and acidity. *B. coagulans* possess a wide range of probiotic attributes such as promoting intestinal digestion, modulating the immune system, preventing pathogenic bacteria colonization, and alleviating certain diseases and gastrointestinal tract (GIT) disorders such as irritable bowel syndrome (IBS), inflammatory bowel disease (IBD) and colorectal cancer [8,20,21]. The predominance of *B. coagulans* in these products was consistent with the claims made by the product manufacturer. Nonetheless, some probiotic species labelled by the manufacturers were not detectable, likely due to the detection limit of our sequencing approach, or the unfavourable storage conditions. *Lactobacillus nagelii*, a common *Lactobacillus* species found in kombucha predominated product H. The species of *Lactobacillus* (LAB) are known to enhance the health-promoting properties of kombucha and widely regarded as probiotics as they can produce short-chain fatty acids (SCFAs), amines, bacteriocins and vitamins involved in antimicrobial and antioxidant activities as well as gut health maintenance, among others [10,12].

To evaluate the health benefits of kombucha, we analysed the functional potential of the microbes across the nine products. The top five out of 59 functional pathways detected were related to sugar metabolism (pentose phosphate pathway (non-oxidative branch), aerobic respiration I (cytochrome c), aerobic respiration II (cytochrome c) (yeast)) and the production of purines (guanosine deoxyribonucleotides de novo biosynthesis II and adenosine deoxyribonucleotides de novo biosynthesis II). Some of the products of these pathways such as ribose-5-phosphate sugar, nicotinamide adenine dinucleotide phosphate (NADPH), adenosine triphosphate (ATP) and purines act as essential growth factors and energy sources for the microbes to synthesise bioactive compounds in the kombucha culture [1].

We further selected 20 functional pathways including SCFAs production, biosynthesis of precursors for B vitamins, amino acids and organic acids [1,4,8]. We noted that functional pathways such as “fatty acid and beta-oxidation II (peroxisome)” and “fatty acid and beta-oxidation I” were only contributed by LAB but not AAB. These pathways metabolised fatty acids to produce acetyl-CoA in an anaerobic condition. Acetyl-CoA is a precursor for the biosynthesis of acetate, one of the common SCFAs that play an essential role in maintaining gut health, immune homeostasis and preventing disease development [22]. LAB, which are facultative anaerobes may utilize these pathways to generate SCFAs when oxygen level is depleted during kombucha fermentation, unlike AAB, which requires oxygen to grow [6,20,23–24].

In comparison, AAB contributed 20% more pathways involved in sugar metabolism, respiration, biosynthesis of amino acids and B vitamins than LAB. For instance, superpathway of aromatic amino acid biosynthesis, superpathway of acetyl-CoA biosynthesis, chorismate biosynthesis I, aerobic respiration II (cytochrome c) (yeast), aerobic respiration I (cytochrome c) and biosynthesis of adenosylcobalamin (vitamin B12) were among the pathways found in AAB but absent in LAB. Meanwhile, lower abundances of species from the “others” group contributed to the 20 selected pathways as compared to AAB and LAB. This could suggest that AAB and LAB possess greater capability in producing beneficial metabolites as compared to non-AAB and -LAB species.

Nevertheless, the exact metabolite composition in kombucha would require further targeted chemical assays. One of the limitations of our study is the small sample size, contributed mainly by the relatively limited number of kombucha brands in Singapore. Future work to assess the stability of kombucha microbiome over different production batches is warranted to understand the consistency of microbial and chemical compositions in kombucha. Clinical studies including *in vivo* tests and bioavailability assessments are indispensable to provide clinical evidence of kombucha’s biological activities in humans, thereby confirming the positive health implications of this popular beverage.

## Conclusion

*Komagataeibacter* species, namely *Komagateibacter saccharivorans*, *Komagataeibacter xylinus* and *Komagataeibacter rhaeticus* were the most prevalent and dominant bacteria present in AAB-dominant products (B, D, G and I) whereas *Bacillus coagulans* and *Lactobacillus nagelii* were the dominant species detected in LAB-dominant products (E, F and H). Although the functional pathways contributed by AAB differed from those contributed by LAB, both categories of bacteria can generate specific organic acids, amino acids, vitamin B compounds and SCFAs with health-enhancing properties in kombucha. Future work to characterise the microbial and metabolite concentrations and their consistencies in kombucha across production batches is necessary.

## Author Contributions

Conceptualization, GT, TKY and JL.; Methodology, JD, HSL, CCW, JLWJ; Writing – Original Draft Preparation, JD, HSL.; Writing – Review & Editing, GT, TKY, JL, CCW, JD, HSL. All authors have read and agreed to the published version of the manuscript.

